# KeySDL: Sparse Dictionary Learning for Keystone Microbe Identification

**DOI:** 10.1101/2025.08.07.669165

**Authors:** Max Gordon, Turgut Yigit Akyol, B Amos, Stig U. Andersen, Cranos Williams

## Abstract

Identification of microbes with large impacts on their microbial communities, known as keystone microbes, is a topic of long-standing interest in microbiome research. However, many approaches to identify keystone microbes are limited by the inherent nonlinearity and state-dependence of microbial dynamics. Machine learning approaches have been applied to address these shortcomings but often require more data than is available for a given microbial system. We propose a keystone identification approach called KeySDL which reduces the amount of data required by incorporating assumptions about the type of microbial dynamics present in the experimental system. The data are modeled as originating from a Generalized Lotka-Volterra (GLV) model, an architecture commonly used to simulate microbial systems. The parameters of this model are then estimated using Sparse Dictionary Learning (SDL) Compared to existing methods, this approach allows accurate prediction of keystone microbes from small numbers of samples and provides an output interpretable as reconstructed system dynamics. We also propose a self-consistency score to help evaluate whether the assumption of GLV dynamics is reasonable for a given dataset, either through the application of KeySDL or other analysis tools validated using GLV simulation.

## 1 Introduction

In ecology, keystone species are generally considered to be those with a disproportionate impact on the interaction structure and stability of the ecosystem when removed [1–3]. Identification of keystone species in microbial systems is often sought to help explain microbial behavior or identify means to modulate microbial interactions [4, 5]. In this work we will consider the keystone microbes of a system to be those whose absence has the largest impact on the observed steady state composition of the microbial ecosystem. Identifying this type of keystone species by iterative removal from a microbial community is not generally possible in natural microbiomes with current experimental methods [6]. In synthetic communities it is possible to omit strains from the inoculant used [7, 8] but this very quickly becomes experimentally infeasible to explore as the size of the community increases and with it the required number of dropout screens.

High throughput amplicon sequencing data presents an opportunity to quantify the relative abundance of individual microbes at scale and use the community structure of these broad datasets as a basis for hypothesis generation [5]. However, despite the potential of these data, the analytical tools to transform them into a comprehensive understanding of the structure of a microbial community are still lacking.

One approach that has seen recent use in attempts to identify keystone microbes without individually removing strains is the construction of correlation or cooccurrence networks [4, 9–13]. These networks can be very interpretable. However, they cannot capture the nonlinearity of microbial dynamics. Since this nonlinearity arises in part from the combination of growth and interactions, this prevents these networks from providing a comprehensive view of the underlying microbial system. They are also unable to represent the state-dependency of many microbial interactions, where the behavior of and interactions between microbes in the system depends on the composition of the community as a whole. Co-occurrence networks may provide an overview of a particular state of the microbiome but will not reliably result in insights into perturbations from that state. Additionally, the edges in these networks do not necessarily correspond to direct interactions between microbes as they capture associations, which may also be present between microbes with shared influencing factors. Although correlation network approaches are widely used, their sensitivity is limited [14]. Prior tests attempting network reconstruction from simulated data have shown that co-occurrence networks cannot reliably recover keystone species due to these limitations [11, 14].

Machine learning (ML) approaches have been applied with some success to the problem of keystone microbe identification [6, 15, 16]. The Data-driven Keystone species Identification (DKI) framework based on cNODE2 uses a compositional neural Ordinary Differential Equation (ODE) to implicitly infer microbiome assembly rules from presence/absence vectors representing species present in a given sample [6]. This approach functioned reliably in simulation and with a small in vitro synthetic community. However, this test was performed with relatively large numbers of samples and small numbers of microbes compared to more typical experiments. The models produced by this framework are multi-layered neural networks that may be difficult to interpret for further insights beyond keystoneness predictions. The number of observations required to constrain an ML model will grow with its complexity and number of parameters [17]. This can pose a major challenge for microbial studies, where large numbers of observations are often not experimentally feasible, so it is often desirable to use a simple model with fewer parameters when limited data is available.

Another approach is the inference of models of microbiome dynamics, often using Generalized Lotka-Volterra (GLV) models [18–20]. These approaches typically require time-series datasets and absolute microbial abundances, rather than the compositional measurements typically available. When appropriate data is available and the scale of the system permits, interpretation of these structures can be very insightful and connected to relevant biological concepts such as growth rates.

Amplicon sequencing measurements of microbial systems are compositional [21]. As a result, an analog of the GLV system that can operate on compositions is desirable for any analysis of an assumed-GLV system through compositional observations. Replicator dynamics constitute such a compositional analog to the GLV model capable of capturing the behavior of the composition of a GLV system without modeling its total abundance [22].

We propose an approach called Keystone Sparse Dictionary Learning (KeySDL) to predict microbial keystoneness based on the assumption that the system we are observing is a Generalized Lotka-Volterra system under steady state. To do this, we first explore an operational definition of keystoneness based on Bray-Curtis dissimilarity of relative abundance profiles in the context of a GLV system. The assumption that the system is under steady state removes the need for numerical integration of the system dynamics, simplifying model optimization. We then solve for the parameters of this GLV system or the corresponding compositional replicator system using Sparse Dictionary Learning (SDL) [23] with additional constraints based on system traits typically observed in experimental microbial communities such as finite carrying capacity. These additional constraints allow our model to converge to a solution from a very small number of observed system steady states, making KeySDL a viable option for datasets where GLV dynamics are a reasonable assumption and there is not enough data available to constrain more flexible ML models. In addition to estimates of impact on removal, KeySDL estimates the parameters of the GLV or replicator system that best describes the observations, providing an interpretable view of model parameters. We also develop a self consistency measure to evaluate the potential validity of assuming GLV dynamics govern a dataset. Beyond keystone identification, the self consistency measure may be useful in identifying scenarios where GLV simulations are not an appropriate in silico test for analytical methods.

## 2 Methods

### 2.1 Steady States and Keystoneness in GLV Systems

In this work, we consider a microbial system governed by Generalized Lotka-Volterra dynamics. This approach does have limitations in the complexity of microbial interactions it is capable of representing but has nonetheless found widespread use in simulation [6, 15, 16, 24, 25] and analysis [18–20] of microbial systems. The fixed growth rates and interactions mean this model is unable to capture changing external influences on the system but this is a shortcoming shared with many other approaches, including nonlinear machine learning methods.

We will consider an *n*-microbe GLV system with set of species 𝕊 = 1, 2, …, *n* given by:

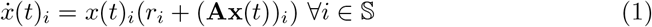

with **x**(*t*) ∈ ℝ^*n×*1^ representing microbial abundances over time, **r** ∈ ℝ^*n×*1^ microbial growth rates, and **A** ∈ ℝ^*n×n*^ microbe-microbe interactions. This system has several steady states denoted 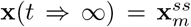, where *m* is an index associated with each steady state of the system. The trivial steady state is achieved when:

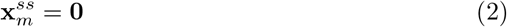

and corresponds to the extinction of all species. At the steady state:

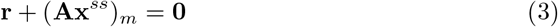

all microbes are present in some quantity given by 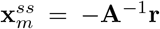. Finally, for all remaining 2^*n*^ − 2 steady states, we consider combinations of extinct microbes 𝔼 and non-extinct microbes ℕ, such that 𝕊 = 𝔼 ∪ ℕ and 𝔼 ∩ ℕ = ∅. This additionally means that we can associate each steady state index *m* with a species presence profile ℕ and extinction profile 𝔼. For every possible 𝔼 and ℕ there exists a steady state where:

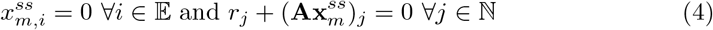

In this context, the keystoneness of a given microbe is the distance, by any keystoneness metric, between the steady state with no extinctions and a perturbed steady state where the microbe is extinct.

It is possible for any steady state in a GLV system to require negative abundances. However, this is not achievable in a real microbial system nor an ideal GLV system as the GLV equation does not allow zero crossing. Therefore, in the case that a perturbation places the microbial community in a steady state where some microbes are required to be negative, these microbes will be driven to extinction since their abundances cannot be negative. If the steady state corresponding to this expanded set of extinct microbes also has microbes with negative abundances, the process will repeat until the system reaches a steady state with all non-negative abundances. This corresponds neatly with the definition of keystoneness by the decrease in the number of distinct species observed in a system after removal of the target microbe. Despite this symmetry, we use Bray-Curtis dissimilarity to measure keystoneness to capture changes that fall short of complete species extinction. In order to avoid simply measuring the abundance of a microbe, we also exclude the microbe’s own abundance prior to the calculation. This can be expressed by defining 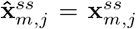 where *j ≠ i*, and 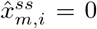, setting the microbe *i* to 0 abundance. The Bray-Curtis Keystoneness of microbe *i* is then as follows:

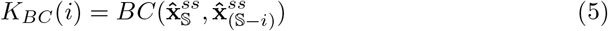

where *BC* is Bray-Curtis dissimilarity [26] and 𝕊 is the set of all microbes in the modeled system.

Because the steady states of a GLV system are defined by the combinations of extinct and nonextinct strains, there is only one steady state for each combination of microbes, resulting in a total of 2^*n*^ steady states. This means that for a system following GLV dynamics we should not expect perturbations falling short of complete removal of a species to cause any change in the steady state the system will return to. As a result, we focus in this work on keystoneness defined by microbial dropouts. This also implies that a microbial system where changes in species abundance without extinction or addition of new species lead to changes in steady state must be governed by non-GLV dynamics such as stochastic extinction.

### 2.2 KeySDL Recovery of GLV Model Parameters

To evaluate keystoneness, our goal is to recover a model with the same steady states as the original system. By focusing only on steady states, we can ignore the nonlinearity of GLV dynamics and solve Eq. 1 at 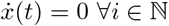 to find **A** and **r** such that:

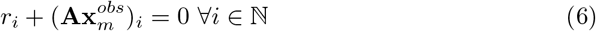

for each observation 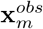with non-extinct microbes ℕ. We assume that each observation is a measurement of an underlying system steady state, 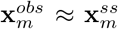. For any reasonable number of observed steady states, this creates an underdetermined system of linear equations, which may be solved as a Sparse Dictionary Learning (SDL) problem [23]. In SDL, underdetermined systems of linear equations may be solved if we have prior knowledge that the solution will be sparse, which further constrains the set of possible solutions. By imposing sparsity through penalty or learning structure, we can approximately solve a system that would otherwise not have a single solution [23].

There are many possible ways to enforce sparsity, but in many practical cases L1 regularization is optimal or nearly so [27]. In the case of learning GLV model parameters, there are several additional constraints that can be incorporated based on knowledge of the underlying system. These are 1) **A** must be invertible for a steady state to exist and 2) All diagonal entries of **A** must be negative. This is because the diagonal entries of **A** represent the interaction between a microbe and itself. If any of these values are positive, unbounded growth would be possible as more of a microbe being present would further increase its growth rate. This would correspond to a microbial system which would not exist in reality.

To allow flexible inclusion of these constraints as well as extension to other constraints in the future, we solved this optimization problem with gradient descent in Torch [28]. Sparsity was enforced by an L1 regularization on the entirety of **A**, while invertibility was enforced with a penalty for a small determinant of **A** and the negative diagonal was enforced with a large negative L1 penalty on diagonal entries of **A** larger than −0.1. The L1 regularization requires selection of a regularization weight, which may be accomplished by cross-validation for each individual dataset. However, in this work we selected a value of 1 × 10^*−*15^ based on the simulation testing and carried this parameter through to the experimental datasets. This is a relatively small value for an L1 regularization because the scale of the error is small relative to the number of parameters subject to the L1 penalty. Higher L1 values will often result in the regularization contributing more to the objective function than the error, leading to a prediction of **A** = **0**_*n,n*_. The weight of the penalty terms for invertibility and diagonal negativity are in theory tunable parameters but in practice they only need to be sufficiently large that they outweigh any improvement in reconstruction error that might come from violating them. A graphical overview of the KeySDL approach applied to a GLV system is shown in Fig. 1.

**Fig. 1:**
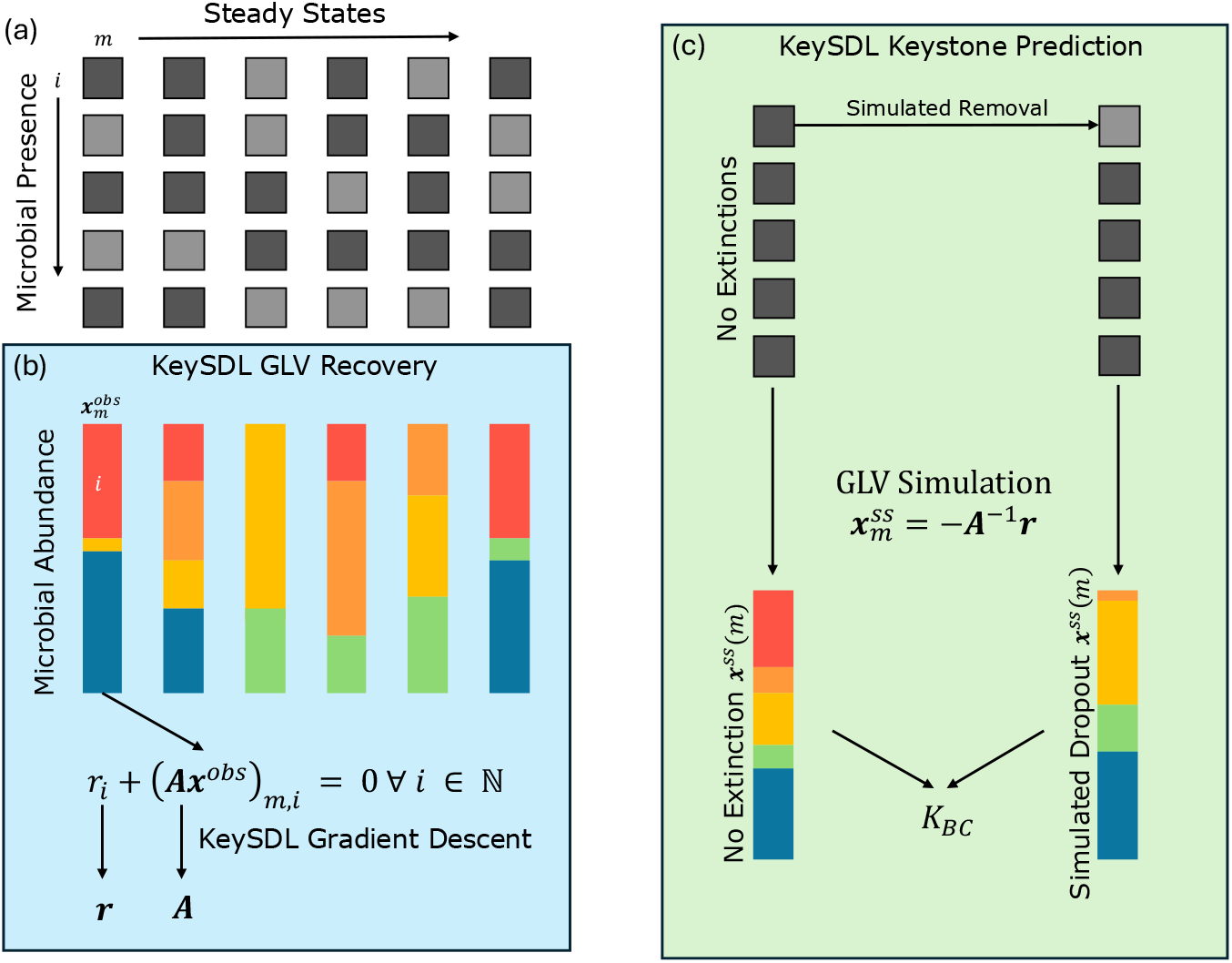
Key concepts in the KeySDL pipeline applied to a GLV system. (a) The presence and absence of each microbe *i* defines steady states *m*. (b) Each measured steady state has microbial composition 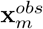. These observations are used in combination with Eq. 6 to solve for interactions **A** and growth rates **r**. (c) With estimates of these model parameters, we can predict compositions for both simulated no-extinction and simulated single-extinction systems for each microbe *i* to derive an estimate of impact on removal *K*_*BC*_(*i*).

### 2.3 Relating GLV and Compositional Replicator Dynamics

While GLV models operate on absolute abundances, high throughput sequencing measurements are inherently compositional [21]. Without measurement of the total quantities present, these sequencing data only convey the ratios of microbes in the system to each other. Attempting to completely model a GLV system based on compositional measurements is not possible due to the loss of information compared to the full abundance profiles.

Replicator systems are a way of modeling compositional measurements of dynamics, following the form:

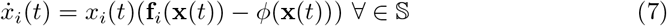

Where *f* (**x**)(*t*) is a function describing the fitness of element x over time and *ϕ*(**x**) = Σ_*j*_ *x*_*j*_*f*_*j*_(**x**(*t*)). In the case of the fitness function **f** (**x**(*t*)) = **Fx**(*t*) it is possible to map trajectories from an *n* dimensional GLV system to an *n* + 1 dimensional replicator system [22, 29]. An additional dimension is needed to encode the system’s total abundance. Without this, there is not full equivalence to the GLV system but trajectories in the GLV and replicator systems do have equivalent compositions [29]. As a result, the relative abundance profiles of the steady states of the replicator system will be the same as the relative abundance profiles of the steady states of the GLV system. This allows us to use replicator dynamics to model steady states and keystoneness from a compositional view of an assumed GLV system.

This replicator system does not directly model the growth rates **r** of the original GLV dynamics but the impact of the growth rates on the composition is in the interactions matrix **F** by the relationship 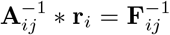 [29]. The impact of the growth rates on the true abundances is outside the scope of modeling for a replicator system.

Finally, steady states of the replicator system are difficult to find compared to those of a GLV model. We can use the equivalence of GLV and replicator systems to find these steady states without numerical methods. A GLV model with interactions matrix **F** and uniform growth rates **r** = [1, …, 1] will have the same steady state compositions as a replicator model with interactions matrix **F** [29]. This allows for rapid computation of the steady state relative abundance a set of microbes will converge on under replicator dynamics without full simulation.

The framework used to fit GLV dynamics in Section 2.2 is easily extended to compositional data using the equivalence of GLV and Replicator dynamics [22, 29]. Non-zero steady states of a Replicator system occur when:

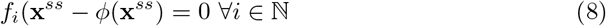

We can substitute this into the KeySDL optimization procedure and recover the **F** matrix of the Replicator system associated with the composition of the underlying GLV dynamics, which is related to the steady states and therefore keystones as described earlier.

### 2.4 Evaluation of Steady State Consistency

One concern with the application of a model that assumes GLV dynamics is whether GLV dynamics are a reasonable approximation of the underlying system. We propose a self-consistency metric *S*_*sc*_ which evaluates how well GLV dynamics are able to capture the system’s behavior to provide insight into the suitability of GLV dynamics for modeling an individual dataset. This score is a measure of how well the steady states of the reconstructed system correspond to measured experimental samples. *S*_*sc*_ is constructed by evaluating the Bray-Curtis dissimilarity between the predicted steady states and true samples for each sample in the training dataset. The mean of these dissimilarity measurements is subtracted from 1 to create a self-consistency metric that is 1 if the data is perfectly explained by a GLV model. Given *N* observed steady states 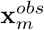 and reconstructed steady states 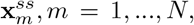, we can compute the Bray-Curtis dissimilarity between each pair as 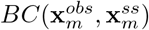 and then assemble the self-consistency metric as:

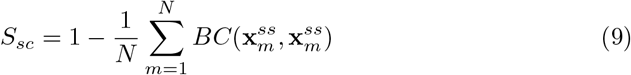

### 2.5 Cross-Validated Prediction for Single-Dropout Datasets

To validate KeySDL, we need datasets with ground-truth *K*_*BC*_ values. The most available type of experiment where this can be computed are those consisting of dropouts of individual microbial strains. One challenge arising from this experimental structure is that we cannot use KeySDL on all the data at once without the model seeing the dropout sample it is trying to recreate during training. We use 3-fold cross-validation for prediction from single dropout datasets to address this. In this scheme, the data are divided into 3 folds. These are split into 3 training sets each consisting of 2 folds, and KeySDL is used on each training set. Predictions are made for the microbes whose dropout samples are not part of the training set. By combining these predictions from the 3 folds, each microbe’s predicted keystoneness is based on a model trained without seeing the dropout it is predicting.

This approach is conceptually very similar to leave-one-out cross-validation. We selected 3-fold instead as with an *N* sample dataset it trains the models on 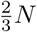 samples rather than *N* − 1 samples. This provides a more rigorous test of KeySDL analogous to other datasets where dropouts for each individual strain are not available.

## 3 Results

### 3.1 GLV Reconstruction with KeySDL from In-Silico Absolute Abundance Data

To validate KeySDL’s ability to reconstruct GLV dynamics from absolute abundance data, we generated ground-truth GLV models and simulated different dropout combinations to produce a synthetic dataset from each set of parameters. We generated the **A** matrices using the Klemm algorithm [30], which produces small-world scale-free networks, properties consistent with interactions in observed ecosystems, while **r** was drawn from a uniform distribution between 0.1 and 1.

First, we generated a 50-microbe GLV system and evaluated the KeySDL reconstruction accuracy based on 500 randomly generated multiple dropouts. The total number of steady states for this system is 2^50^, so 500 is a small sample of the total number of steady states. The ground truth and reconstructed dynamics are very similar, with a mean squared error of 9.5*e* − 4 for **A** and 5.5*e* − 3 for **r**. Qualitatively, the true and reconstructed **A** matrices are shown in Fig. 2a and show the same overall structure. True and reconstructed *K*_*BC*_ values are shown in Fig.2b. The reconstruction perfectly captures the rank order of keystones (*ρ* = 1, *p* = 7.3*e* − 91) and closely reproduces the GLV interactions matrix. As a basis for comparison with existing cooccurrence network generation approaches[14], we generated a co-occurrence network using Pearson correlation from the simulated absolute abundance data and computed network degree and betweenness centrality, shown in Fig. S1a and S1b. Both degree (p=0.12) and betweenness (p=0.76) are uncorrelated with the removal impact *K*_*BC*_, demonstrating that these network topological metrics do not provide a useful way to assess potential impact on removal in this case.

**Fig. 2:**
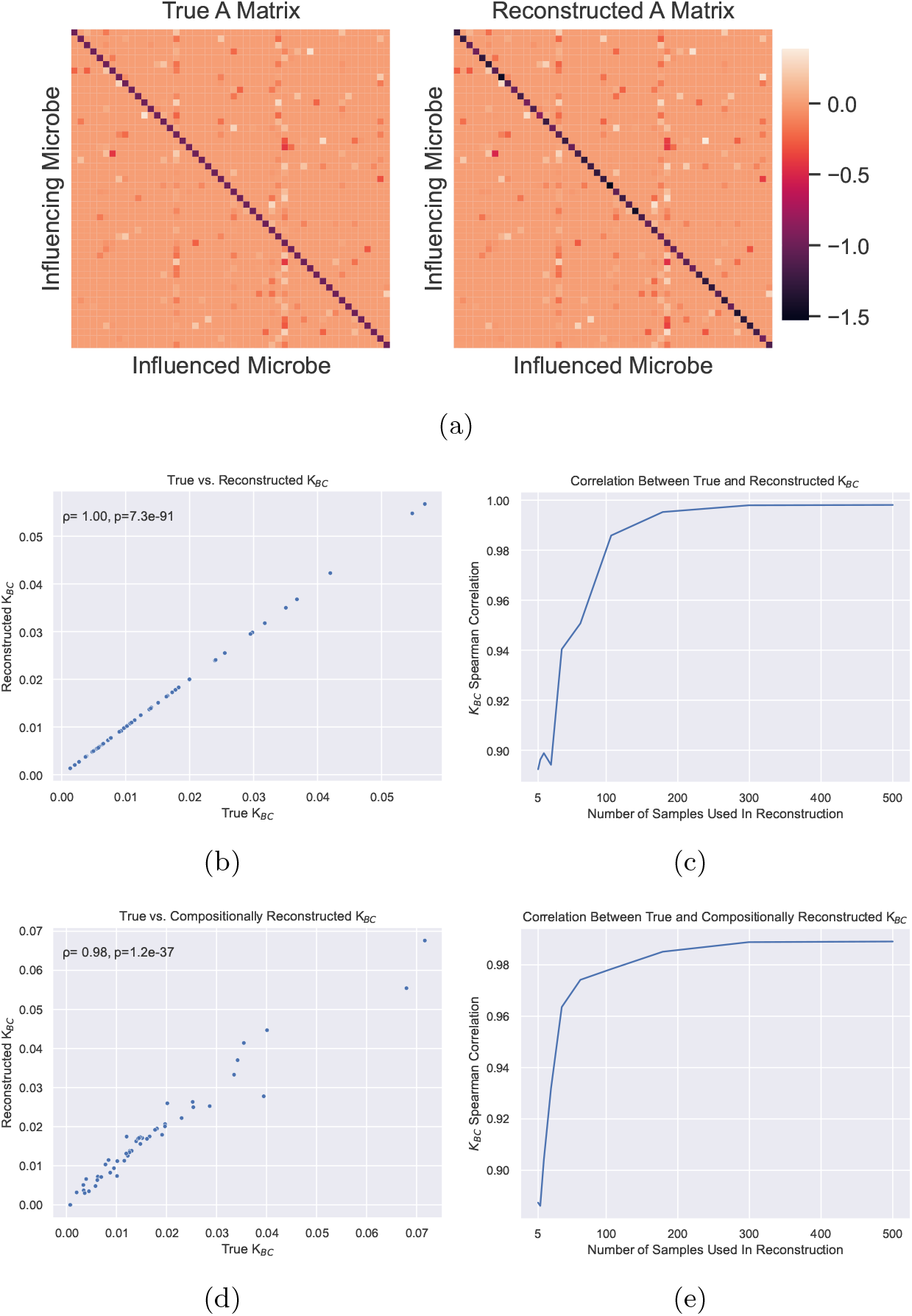
2a) True and reconstructed interactions matrices from 500 samples taken from a GLV model. 2b) True and reconstructed *K*_*BC*_ from 500 samples taken from a GLV model. 2c) Spearman correlation between true and reconstructed *K*_*BC*_ with varying numbers of samples from a GLV model (*ρ* = 0.1, *p* = 7.3*e* − 91). 2d) True and compositionally reconstructed *K*_*BC*_ from 500 samples taken from a GLV model. 2e) Spearman correlation between true and compositionally reconstructed *K*_*BC*_ with varying numbers of samples from a GLV model (*ρ* = 0.98, *p* = 1.2*e* − 37).

Next, we also evaluated how the number of distinct steady states used to create the reconstruction impacted reconstruction accuracy. This was done by selecting a subset of multiple dropouts for each number of dropouts being tested and computing the average spearman correlation between true and reconstructed *K*_*BC*_ based on individual strain dropouts. This was repeated for 10 random subsets to prevent any individual sampling from distorting the results. We tested a with numbers of observed steady states ranging from 5 to 500. The average *K*_*BC*_ is shown in Figure 2c. This correlation is above 0.9 whenever more than 20 steady states are used in the reconstruction and remains above 0.85 even at 5 steady states, indicating that only a small number of observations are needed to usefully constrain the reconstruction.

### 3.2 Replicator Reconstruction with KeySDL from In-Silico Relative Abundance Data

With KeySDL tested on absolute simulated data, we next validated its performance on compositional data like those produced by real microbiome amplicon sequencing experiments by repeating the same testing steps with compositionally transformed data. Notably, the interactions matrix recovered by compositional KeySDL is not the same as the original GLV interactions matrix but is instead a reconstruction of the Replicator interactions matrix associated with its composition. This relationship is given by 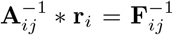. We demonstrate the correlation (*ρ* = 0.98, *p* = 1.2*e* − 37) between true and reconstructed *K*_*BC*_ for a compositional reconstruction from 500 samples in Fig. 2d. Degree and betweenness for a co-occurrence network produced from the simulated compositional data using SparCC [12] are shown in Figs. S2a and S2b. These network metrics are not correlated with the true impact on removal and would therefore not be useful in identifying keystone microbes. Finally, the impact on reconstruction of keystoneness from numbers of steady state observations ranging from 5 to 500 is shown in Fig. 2e. As with the GLV reconstruction from absolute abundances, the replicator reconstruction from relative abundances is able to accurately estimate *K*_*BC*_ for all microbes in a system from a small number of samples.

### 3.3 Evaluation of KeySDL Robustness to Noise

So far, we have evaluated the model with noiseless samples of different steady states. However, real data will contain read noise and will be quantified into read counts prior to normalization, which introduces additional noise. To evaluate the robustness of KeySDL reconstruction to noise, we added Gaussian noise with standard deviation varying from 0% to 200% of the mean microbial abundance into 200 samples used to estimate a replicator system with KeySDL and evaluated its ability to recover *K*_*BC*_ as noise increased. This was performed 5 independent times for each noise level and the results averaged. These results are shown in Fig. 3a and demonstrate that the reconstruction is able to recover keystones even in the presence of large amounts of noise, although with reduced accuracy, maintaining a Spearman correlation between true and predicted keystones above 0.7 until more than 100% of the mean microbial abundance is added as noise.

**Fig. 3:**
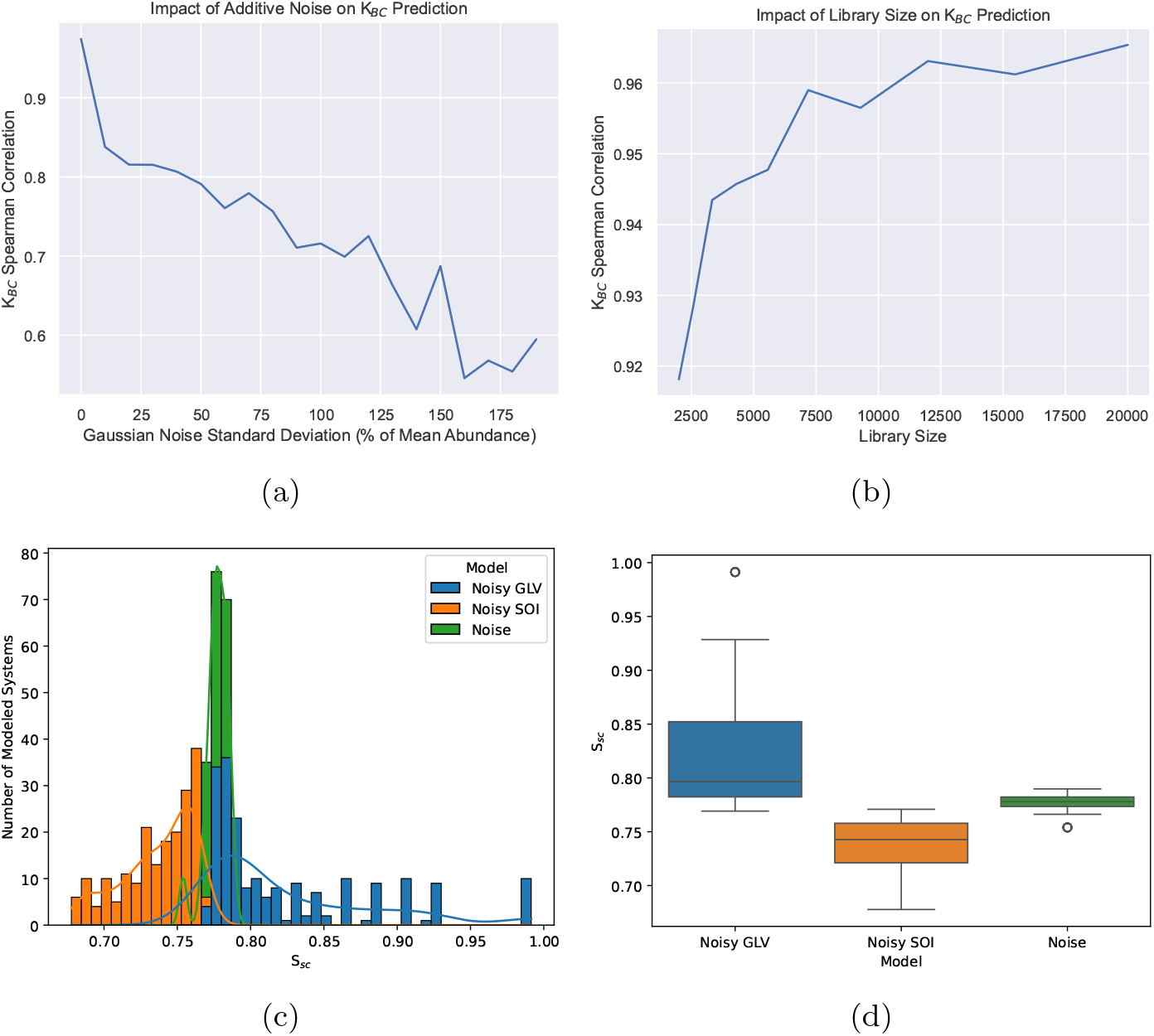
3a) Spearman correlation between true and compositionally reconstructed *K*_*BC*_ with varying Gaussian noise added to 100 samples from a GLV model. 3b) Spearman correlation between and between true and compositionally reconstructed *K*_*BC*_ from 100 samples from a GLV model with different library size. 3c) Histogram and 3d) boxplot of consistency score *S*_*sc*_ for SOI-simulated data, GLV-simulated data, and random noise. The different model specifications have different median values (*p* = 1.35*e* − 99)

The total number of reads collected, library size, is an important property of amplicon sequencing datasets. Since all abundances must be expressed as a fraction of the reads collected and fractional reads are not possible, a small library size corresponds to quantization noise in the abundance measurements. We quantized the data targeting a total library size across the entire sample to evaluate KeySDL’s robustness to this experimental variable. If the library size is decreased, each quantization step will be larger, resulting in higher quantization noise. This was performed 5 independent times for each library size and the results averaged. The results of this test are shown in Fig. 3b and demonstrate the reconstruction’s robustness to sampling and quantization, even with small library sizes, with a Spearman correlation between true and predicted keystones above 0.91 even for the smallest library size tested of 2500 reads.

### 3.4 Characterization of Self-Consistency Scoring

Next, we evaluate how the self-consistency metric *S*_*sc*_ allows us to distinguish between datasets that are consistent with GLV dynamics and those that are likely derived from other system structures. To measure this, we compared *S*_*sc*_ for synthetic datasets generated with GLV dynamics, Self-Organized Instability (SOI) dynamics [31], and pure noise. We used miaSim [25] to produce the SOI simulations. We added Gaussian noise with standard deviation varying from 0 to 200% of the mean microbial abundance to simulate the effect of experimental variation and sequencing noise. This test was performed for 10 synthetic datasets of each model type and noise level, with the aggregate results shown in Figs. 3c and 3d. There is a distinction between synthetic data from GLV and SOI systems, with samples comprised entirely of noise in between. These distributions are distinct, with a Kruskal-Wallis p value of 1.35*e* − 99. From this plot, we can expect that a noisy system reasonably approximated by GLV dynamics will have a self-consistency score of 0.75 or greater. Notably, there is overlap between the score distributions for noise-perturbed GLV systems and measurements entirely comprised of noise. Despite this, a system operating under non-GLV dynamics is still identifiable through *S*_*sc*_. While this means that *S*_*sc*_ in a range consistent with a GLV model is not a conclusive sign that the KeySDL pipeline will produce accurate predictions, it does provide a way to avoid application of the approach in cases where a GLV model is highly misspecified.

### 3.5 Validation of KeySDL In Vitro

To validate the performance of KeySDL in recovering the behavior of experimental biological systems, we identified an in vitro gut microbial SynCom experiment with individual species dropouts [7]. The SynCom in this experiment consists of 14 gut microbes grown in a batch bioreactor. The dropouts in this dataset allow us to compute ground-truth *K*_*BC*_ in addition to fitting a KeySDL reconstruction, enabling direct comparisons between the model’s reconstructions and measured keystones.

In order to predict *K*_*BC*_ for each microbe from a model not trained on that microbe’s dropout, we used the cross-validation scheme described in 2.5. The results of this cross-validation are shown in Fig. 4a, along with degree and betweenness centrality for a co-occurrence network generated using SparCC [12] on all data at once in Fig. S3a and S3b. The KeySDL reconstructed keystones reasonably correspond to the measured keystones of the microbial system (*ρ* = 0.71, *p* = 4.2*e*−3). The co-occurrence network metrics do not have a statistically significant association with measured BrayCurtis keystoneness, despite being fitted to all data at once. The validity of *K*_*BC*_ as a biologically relevant metric is also supported by the recovery of *E. coli* and *B. dorei* as the primary keystones of the system, with predicted *K*_*BC*_ values of 0.37 and 0.22 respectively. These microbes were identified as dominant microbes in the system by the authors of the original paper, and while they are also highly central based on the two network centrality metrics, there are many other microbes with comparably high centrality by those measures. *S*_*sc*_ for this dataset is 0.81, which is within the range expected for a noisy GLV system and suggests that it is reasonable to reconstruct this system based on GLV dynamics. For a qualitative view of this, the measured and reconstructed compositions are shown in Fig. 4c, along with compositions from a null model consisting of removal of the dropped-out microbe followed by re-scaling. Notably, KeySDL is able to capture the increase in *B. adolescentis* when *E. Coli* is removed, as well as the presence of *L. symbosium* in notable quantities in nearly all dropouts.

**Fig. 4:**
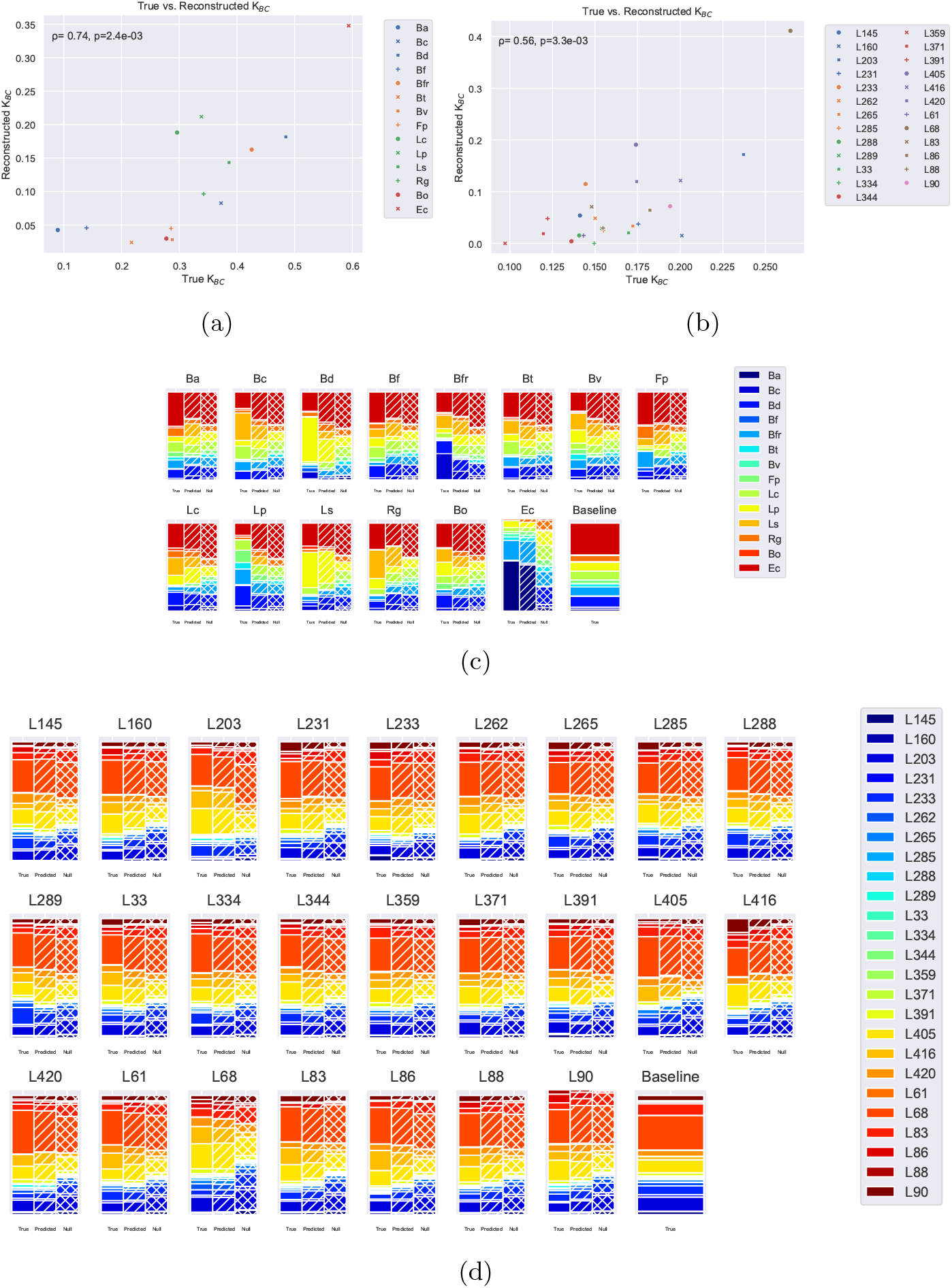
Spearman correlation between predicted and measured *K*_*BC*_ with data from 4a) [7] (*ρ* = 0.74, *p* = 2.4*e* − 3) and 4b) [8] (*ρ* = 0.56, *p* = 3.3*e* − 3). Measured composition, KeySDL reconstruction composition, and null model composition for data from 4c) [7] and 4d) [8]. The null model consists of removing the dropped-out microbe and scaling all other abundances to preserve the sum.

### 3.6 Validation of KeySDL In Planta

In addition to in vitro test data, it is desirable to validate the KeySDL methodology in a more complex system. For this, we identified a plant microbial dataset with a suitable experimental setup. The data used for this validation consists of sequencing of the root microbiome of *Arabidopsis thaliana* grown in a SynCom, with 25 microbes selected for single dropouts, as well as a complete SynCom with no dropouts as a control. There are two biological replicates, with each consisting of 8 technical replicates [8].

We predicted *K*_*BC*_ according to the cross-validation scheme described in Section 2.5. Plotting these predictions against the *K*_*BC*_ derived from the dropout and control samples results in the plot shown in Fig. 4b. KeySDL identified L68, L405, and L203 as top keystones. L68 was identified as the dominant strain in the system in the original paper, while L203 and L405 had significant impacts on community composition when removed. The reconstructed and measured keystones of the system are correlated (*ρ* = 0.56, *p* = 3.3*e* − 3). In addition to the KeySDL reconstruction, we produced cooccurrence networks from the same data using SparCC [12], shown in Figs. S4a and S4b. These cannot be cross-validated, so the co-occurrence network generation uses all the dropouts we are trying to predict the impact of. Despite this, neither co-occurrence network metric is correlated with *K*_*BC*_ for either replicate. Computing *S*_*sc*_ to evaluate the reconstruction of individual steady states, we obtained a score of 0.92. This score is in the range expected for GLV dynamics from in silico testing, so application of KeySDL here is reasonable. The measured and reconstructed compositions that *F*_*sc*_ is derived from are shown in Fig. 4d, along with a null model where the composition is rescaled with the dropped-out microbe removed. In several dropouts, such as L288, L344, L88, and L90, the KeySDL accurately show the presence of L86 where the null model does not.

### 3.7 KeySDL Recovers Disproportionate Keystones In Silico

Proportionality is a major component of the ecological definition of a keystone species [1]. This is the idea that a keystone species must have an impact larger than would be expected based on its abundance. Prior work in microbial keystone identification, however, has largely considered impact on removal independently of starting abundance [11, 32]. Although in these prior works the measure of impact on removal does exclude the removed microbe’s own abundance, to evaluate proportionality we need to compare against a measure of mean abundance. We can randomly sample steady states of the reconstructed GLV or Replicator system and compute mean abundance across these steady states. In Fig. 5a we plot this abundance against reconstructed impact on removal for a simulated microbial system created from a network built using the Klemm algorithm [30]. This network construction approach is prone to generating keystones. In the plot we can observe that KeySDL has reconstructed a system where *K*_*BC*_ and abundance are broadly correlated. However, there are several disproportionate keystones with impacts on removal greater than expected given their abundance. These disproportionate keystones may fit the ecological definition of keystoneness more closely than the microbes with the larger impact on removal but also higher abundance.

**Fig. 5:**
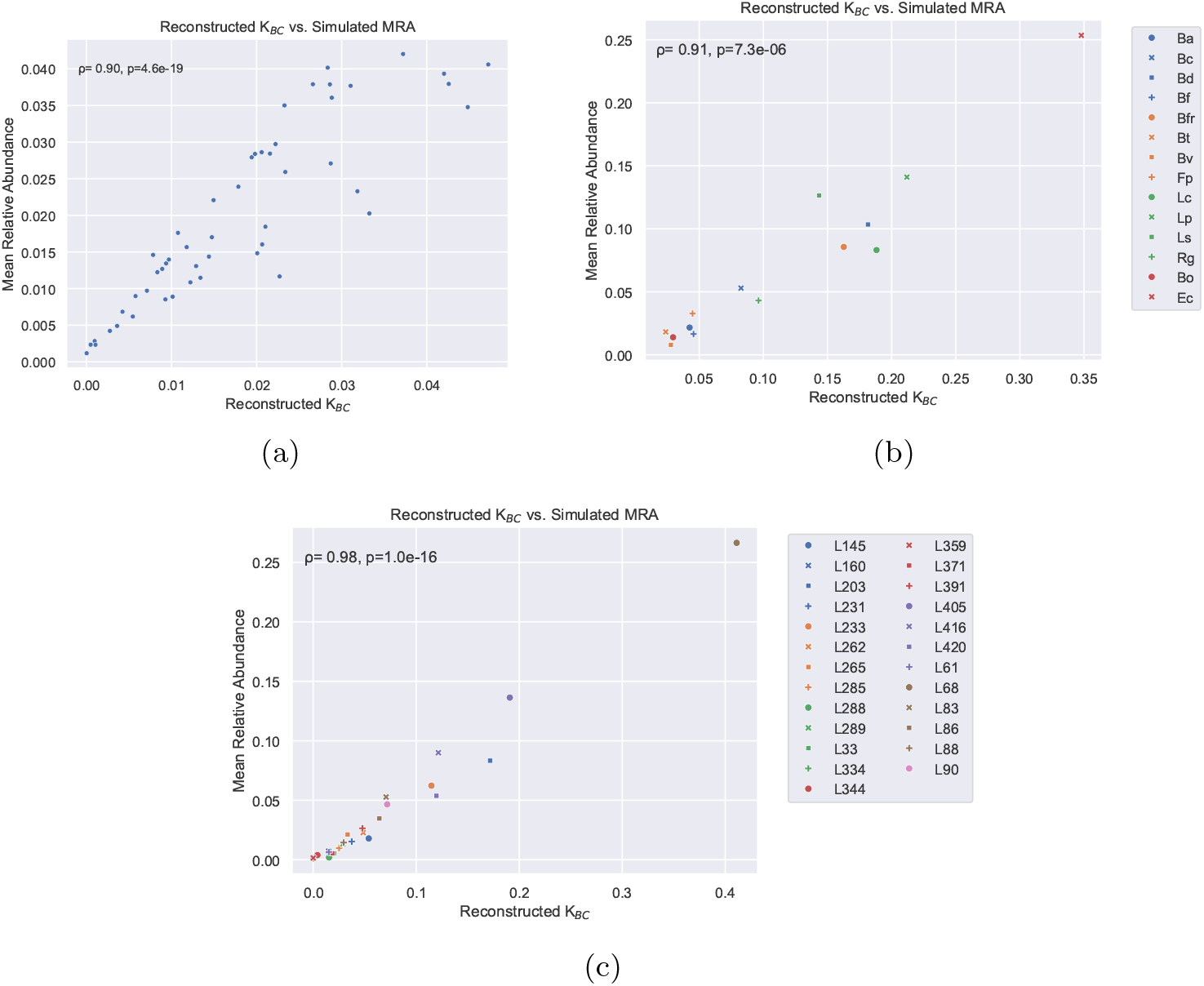
5a) Spearman correlation between reconstructed *K*_*BC*_ and simulated mean relative abundance from 500 simulated steady states reconstructed from a simulated GLV system (*ρ* = 0.90, *p* = 4.6*e*−19). 5b) Spearman correlation between reconstructed *K*_*BC*_ and simulated mean relative abundance from 500 simulated steady states for the dataset from [7] (*ρ* = 0.91, *p* = 7.3*e* − 6). 5c) Spearman correlation between reconstructed *K*_*BC*_ and simulated mean relative abundance from 500 simulated steady states for the dataset from [8] (*ρ* = 0.98, *p* = 1.0*e* − 16).

### 3.8 KeySDL Finds Proportionate Keystoneness in Experimental Data

Following recovery of simulated disproportionate keystones, we also investigated whether keystoneness is proportional to relative abundance in the two validation datasets we identified. We used KeySDL to reconstruct a set of replicator dynamics from each dataset and then extracted impact on removal and mean relative abundance for each microbe from these reconstructions. Plots of these results are given in Figs. 5c and 5b and show that in both cases *K*_*BC*_ and mean relative abundance are correlated without outliers. This suggests that in both of these experimental systems, there may not be strongly disproportional keystones.

## 4 Discussion

We have presented an approach for the reconstruction of microbial dynamics and identification of keystone microbes based on observations of steady states of the microbial system. We demonstrated the robustness of this approach in recovering keystone microbes in silico with both absolute and relative abundance, as well as with the presence of additive Gaussian noise and quantization error from limited read depth. We developed and characterized a self-consistency metric to help identify when an assumption of GLV dynamics is model misspecification. Finally, we validated this method’s performance in keystone identification with in vitro and plant microbial datasets with dropouts. In these experimental datasets, KeySDL recovered top keystone species broadly consistent with the findings of the original publications which produced the data.

This approach is able to function with many methods of defining keystoneness since it directly reconstructs the steady state abundance or composition of the underlying system. Since these reconstructions follow the form of the assumed dynamics rather than a black box estimator, as in some other recent approaches, it may be feasible to perform time-series simulations based on the reconstructed model parameters. This may be of particular interest if quantification of absolute abundances through spike-in or other methods enables reconstruction of a full GLV model rather than the compositional Replicator model.

In addition to simulation of single-dropout experiments, the reconstruction approach within KeySDL allows evaluation of combinatorial effects within the reconstructed system. This could be used to test the keystoneness of combinations of microbes or to compute metrics such as the D1 measure [32], which evaluates keystoneness across a broad sampling of steady states instead of from the no-extinction condition, for datasets where the presence/absence structure of the samples does not allow for direct computation from the experimental data.

A key limitation of the approach we have developed is the strong assumption of GLV dynamics it requires. Although these dynamics are an approximation, we show that for some experimental data, they are informative of the underlying keystones and can robustly represent the observed relative abundances of some experimental systems (Figs. 4c and 4d). In small or noisy datasets, the low number of samples required and robustness to noise of this approach may make it a suitable option despite the limitations of the GLV model. The risk of using KeySDL in cases where it is not reasonable to assume GLV dynamics is mitigated by the self-consistency score.

One particularly important assumption used here is that microbial growth rates and interactions must be constant across the dataset being used. In this initial work, we have attempted to select datasets for which this is the case in order to demonstrate the viability of KeySDL. However, there are many existing datasets, such as Genome Wide Association Studies (GWAS) or cross-sectional measurements, where this assumption is likely not met due to the influence of host genotype or environment. The reconstruction may still find meaningful keystones in such data if this variation is small compared to the strength of the microbial interactions. In order to make full use of commonly available experiments it may be necessary to extend KeySDL to incorporate non-microbial factors with impacts on the microbiome. Without this, we do not have a way to estimate variation in growth rates and interactions across a dataset. Careful interpretation of the biological relevance of identified keystones would therefore be required when non-microbial influences vary across samples. It should be noted that this restriction is not unique to GLV reconstruction approaches and would affect any prediction of microbial keystones or dynamics that does not incorporate the influence of non-microbial variables.

Time-series datasets are somewhat uncommon in microbiome research but there is potential to extend this approach to such experiments using the framework of neural ODEs to allow optimization of the model parameters through a numerical integrator. This concept would be distinguished from prior work using neural ODEs in microbiome research by the use of an assumed dynamic model (GLV/Replicator) and by the addition of constraints such as those used in this work’s reconstruction from steady states. The stiffness of GLV models may lead to numerical stability issues in making such an extension. The neural ODE approach could also be valuable in non-time-series datasets where initial inoculum sequencing is available. These are commonly collected and making use of them to further constrain the reconstruction would provide an avenue to improve the performance of KeySDL.

## Supporting information

Supplementary Informaation

## Code Availability

KeySDL is available at https://github.com/mjgord/KeySDL. The scripts used to produce the results and figures as well as any supplementary information are available at https://github.com/mjgord/KeySDL-Manuscript-Code.

## Acknowledgments

This work was funded by the Novo Nordisk foundation as part of the Collaborative Crop Resiliency Program through the InRoot project (NNF19SA0059362). MG was also funded by the National Science Foundation Graduate Research Fellowship (DGE-1746939) and the NIH/NCSU Molecular Biotechnology Training Program (NIH T32 GM008776).

