## Supplementary Informaation for "KeySDL: Sparse Dictionary Learning for Keystone Microbe Identification"

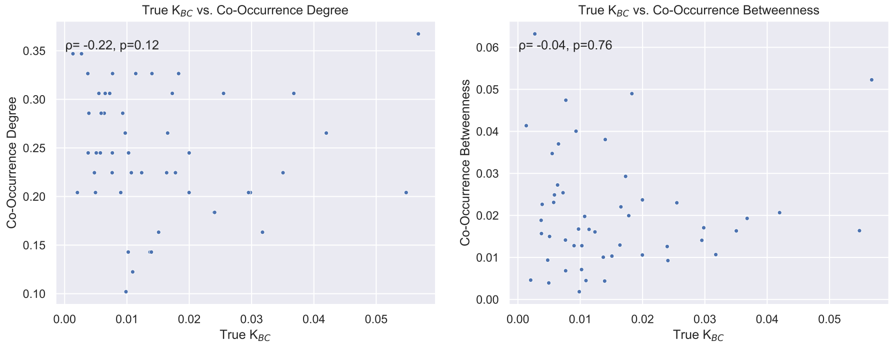


(a) (b)

**Fig. S1**: S1a) Degree and S1b) betweenness of each simulated microbe in a co-occurrence network generated using Pearson correlation.

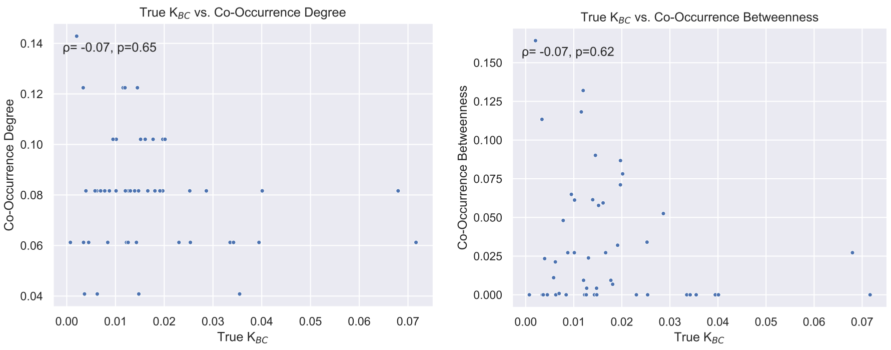


(a) (b)

**Fig. S2**: S2a) Degree and S2b) betweenness of each simulated microbe in a co-occurrence network generated using SparCC [1].

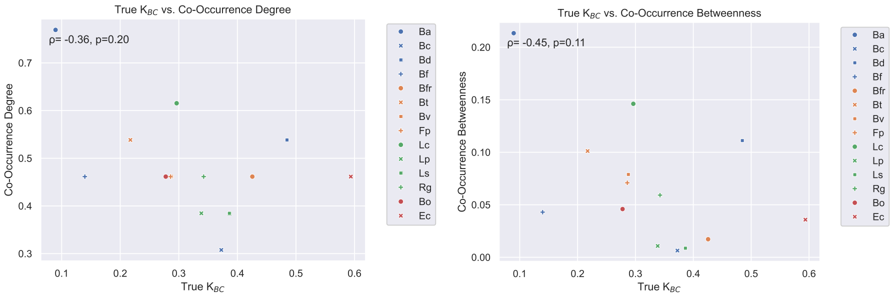


(a) (b)

**Fig. S3**: Lack of Spearman correlation between predicted $K_{BC}$ and S3a) degree and S3b) betweenness centrality from a co-occurrence network produced with SparCC [1] and data from [2].

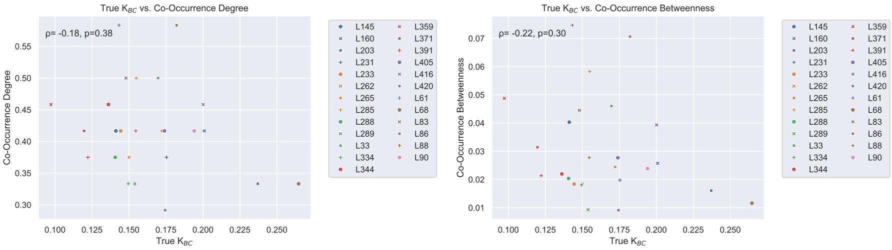


(a) (b)

**Fig. S4**: Lack of Spearman correlation between predicted $K_{BC}$ and S4a) degree and S4b) betweenness centrality from a co-occurrence network produced with SparCC [1] and data from [3].
